# Alpine marmot (*Marmota marmota*) pups emerge increasingly earlier with the ongoing climate change

**DOI:** 10.1101/2025.03.28.645887

**Authors:** Bonenfant Christophe, Cohas Aurélie, Quittet Pierre-Alexandre, Cavailhès Jérôme, Garcia Rébecca, Blache Nicolas, Gaillard Jean-Michel, Douhard Mathieu

**Affiliations:** Laboratoire “Biométrie et Biologie Évolutive” – UMR CNRS 5558, Université Claude Bernard Lyon 1, 43 Boulevard du 11 novembre 1918, Villeurbanne, 69622; CEFE, Univ Montpellier, CNRS, EPHE, IRD, Montpellier, France; Parc National de la Vanoise, 135 Rue Dr Julliand, 73000 Chambéry, France

**Keywords:** Alps, hibernation, phenology, reproduction, Sciuridae

## Abstract

Advance in the phenology of plants and animals is a widely observed response to climate change. The magnitude of the observed changes is, however, very variable across species. Several biological factors could influence the strength of the phenological advances, including lifestyle. Hibernation has evolved in response to harsh environmental conditions and could, hence, potentially buffer organisms against changing climatic conditions. In the Alps, the alpine marmot hibernates for almost 6 months. During that time individuals are sheltered from cold and lack of food, so we could expect alpine marmots to be less responsive to earlier springs than non-hibernating mountain-dwelling species. Here we investigate temporal variation in the date at which pups emerge from their natal burrow for the first time. Using quantile regressions, we provide clear evidence of an earlier pup emergence between 1990 and 2023. Over the study period, the predicted change is of about 4.7 days. In particular, late emergence dates are becoming especially rare over time. Our findings are in line with previous work on other mountain species, which suggests a general advance in reproductive phenology among the organisms living in Alps. The rate of change of pup emergence dates over years is, however, weaker in the alpine marmot than in most other mammalian species studied so far.

## 1 Introduction

Earlier timing of biological events (or phenology *sensu* Morren (1849)) is the core response of living organisms to the ongoing climate change (Parmesan 2006). Tracking the temporal trend of warmer spring temperatures in seasonal environments, plant phenology starts on average almost 3 weeks earlier nowadays than in the 1980s (Badeck et al. 2004, Monahan et al. 2016). Because they rely directly or indirectly on primary productivity, the timing of reproduction, birth, or migration of a wide array of animal species have all been reported to occur increasingly earlier over the last decades (Inouye 2022). A rapid literature survey of studies about the advance of phenology over years in mammalian populations shows an average shift of birth dates ranging from −0.17 days.year^−1^ in yellow-bellied marmots (*Marmotta flaviventris*) to −1.80 days.year^−1^ in North American red squirrel (*Tamiasciurus hudsonicus*; Table 1).

**Table 1:** Observed shifts in birth phenology (measured in number of days per year) of 13 mammalian species recovered from a non-exhaustive literature survey (excluding publications in 2025). The tabulated rates of change are the estimated regression coefficients (*β*) between birth date and time in years extracted from the corresponding reference.

| Species | Scientific name | $\beta$<br>(in days.year<sup>-1</sup>) | Reference |
| --- | --- | --- | --- |
| North American red squirrel | *Tamiasciurus hudsonicus* | -1.80 | Réale et al. (2003) |
| Feral cattle | *Bos taurus* | -1.00 | Burthe et al. (2011) |
| Harbour seal | *Phoca vitulina* | -0.88 | Osinga et al. (2012) |
| Edible dormouse | *Glis glis* | -0.76 | Adamík & Král (2008) |
| Wild dog | *Lycaon pictus* | -0.75 | Abrahms et al. (2022) |
| Wild boar | *Sus scrofa* | -0.55 | Gamelon et al. (2011) |
| Chamois | *Rupicapra pyrenaica* | -0.39 | Kourkgy et al. (2016) |
| Red deer | *Cervus elaphus* | -0.37 | Moyes et al. (2011) |
| Reindeer | *Rangifer tarandus* | -0.29 | Post & Forchhammer (2008) |
| Red deer | *Cervus elaphus* | -0.27 | Bonnet et al. (2019) |
| Roe deer | *Capreolus capreolus* | [-0.16; -0.33] | Hagen et al. (2021) |
| Yellow-bellied marmot | *Marmota flaviventris* | -0.17 | Ozgul et al. (2010) |
| Alpine marmot | *Marmota marmota* | -0.14 | This study |
| Roe deer | *Capreolus capreolus* | -0.08* | Plard et al. (2014) |
| Mean |  | -0.54 |  |

\*this coefficient was not given by Plard et al. (2014) so we re-estimated the quantile regression coefficient (setting $\tau = 0.5$ ) from roe deer data provided in their Appendix S5.

In seasonal environments, birth timing has evolved to match broadly the phenology of food resources, which allows parents to cope better with the increased energy costs of reproduction (Bunnell 1982, Rutberg 1987, Bronson 1989). Climate change is currently disrupting this matching between the timing of food resource flush and of births, the so-called match-mismatch hypothesis (Cushing 1973, Stenseth & Mysterud 2002). The observed advance in phenology of many species is outpaced by the speed of the local weather changes induced by climate change (Visser & Both 2005, Radchuk et al. 2019). This increasing mismatch between the phenology of plants and consumers has profound effects on population dynamics of birds and mammals (Miller-Rushing et al. 2010, Post & Forchhammer 2008, Plard et al. 2014, Rehnus et al. 2020). Also, by its effects on the reproductive success of parents, the differential between the onset of spring and the timing of births also emerges as a major selective force (see Gienapp et al. 2008, for a discussion), with several well documented cases of rapid adaptive responses (Boutin & Lane 2014, Lane et al. 2012).

While the change in reproductive phenology over time is rather pervasive among svertebrate species (Parmesan & Yohe 2003) the direction and magnitude of these observed phenological changes vary a lot among species (Table 1) and across populations (Bronson 2009, Cohen et al. 2018). Contrasting patterns have been documented, even when considering closely related species, making predictions about the effects of climate change on reproductive phenology rather difficult to set. With the accumulation of empirical case studies, some large-scale structuring factors have emerged (Inouye & Wielgolaski 2025). For instance, the ecological consequences of climate changes on plant phenology are particularly strong at high latitudes (Parmesan 2006, Wielgolaski & Inouye 2003, Inouye & Wielgolaski 2025). A similar contrast is expected at low *vs.* high elevation given the usual latitude *vs.* elevation interchangeability in ecology (but see Loewen et al. 2023, for a critique). In western Europe where mild climate is the rule, the ecological consequences of climate change are more frequently reported and of larger magnitude in the high elevation ecosystems of the Alps range than in lowlands (Beniston 2006, Lenoir et al. 2008). High arctic and mountain-dwelling species or populations are therefore of particular concern in face of climate change (Hock et al. 2019).

In addition to environmental factors, the life history and lifestyle of species partly account for the observed variation in the phenological responses of animal species to climate change (Inouye & Wielgolaski 2025). Lifestyle indeed structures many aspects of a species biology (Dobson 2007), including the population responses to climate change. For instance, secondary consumers tend to be less sensitive to climate change than organisms from lower trophic levels due to time lags and the deterioration of the match between food demand and availability at higher trophic levels (Cohen et al. (2018), Mahoney et al. (2020), but see Abrahms et al. (2022) on Wild dogs *Lycaon pictus*). Similarly, hibernation, a rather frequent lifestyle displayed by mammals (Geiser 2013) slows down the pace-of-life of species for a given body mass (Turbill et al. 2011). Incidentally, a slow pace-of-life is associated with more acute effects of climate change (Orgeret et al. 2022).

The predominant ultimate cause for the evolution of hibernation is the prolonged food shortage or harsh seasonal conditions (Davis 1976). The duration and timing of hibernation indeed are the result of a time allocation trade-off between the benefits of a reduced metabolism to hibernate long enough to emerge at a favorable time of the year, and the time needed to build energy reserve during the active season. Despite their adaptation to harsh environmental conditions, hibernating species might be more sensitive to the adverse effects of climate change than non-hibernators (Armitage 2017, Geiser 2013). In marmots, the phenology of reproduction could be particularly affected. Protandry, *i.e.* the tendency for males to emerge before females, is the rule in hibernating sciurids to initiate spermatogenesis (Michener 1984). Thus, males might tracks spring phenology to be able to monopolize and fertilize emerging females. Moreover, most pups of hibernating species are altricial, so their dates of birth and emergence are crucial because a too late emergence might jeopardize individual survival to the next hibernation (Armitage 2014). The high degree of adaptation of marmots to their environment and the constrained life cycle within a year could lead to stronger responses to climate changes of hibernating vs. non-hibernating species (Geiser 2013).

The alpine marmot (*Marmota marmota*) is a fossorial and communal hibernating species (Armitage 2014). For almost 6 months, alpine marmots live underground, relying on body fat accumulated during the vegetative season. Despite this long hibernation, climate-change responses of alpine marmots have been documented on body size (Canale et al. 2016), litter size (Tafani et al. 2013) and juvenile survival (ézouki et al. 2016). All these changes are detrimental to the fitness of individuals (ézouki et al. 2016) and point toward a degradation of the environmental conditions of alpine marmots over the last decades. Like for most hibernating sciurids, the birth of alpine marmot offspring takes place in the burrows and is unobservable directly (Wilson et al. 2016). As a proxy of alpine marmot reproductive phenology, we therefore investigated the temporal change in emergence date of pups, which corresponds to weaning (about 40 days after birth). Alike most of Europe, spring in the Alps is increasingly earlier over the years (Menzel et al. 2006), from which alpine marmots seem to benefit (ézouki et al. 2016). According to the hypothesis of an earlier-spring tracking by herbivores (Inouye 2022), we predicted an advance in the annual emergence date of pups over the years as previously observed in many mammals and birds (Badeck et al. (2004), Monahan et al. (2016); Table 1). With a generation time close to 7 years (ézouki 2018), the alpine marmot has a rather slow pace-of-life for its body size. This slow pace-of-life and the hibernation are two traits that theoretically should make this species particularly sensitive to climate change and the advance of plant phenology (Orgeret et al. 2022, Geiser 2013). We hence expected a higher response of pup emergence phenology for the alpine marmot compared to other mammals.

## 2 Material and Methods

### 2.1 Life cycle of the alpine marmot

Alpine marmots live in families including between 2 and 24 individuals composed of a dominant couple, several sexually mature (≥2 years) and immature (1-year old) subordinates of both sexes, and pups of the year. Each family occupies a territory with a main burrow and a multitude of secondary burrows. The main burrow is made up of a large chamber connected to the above-ground by several galleries. This burrow is used by the family for hibernation and birth. Secondary burrows are mostly used to hide from an immediate danger such as a predator. Being monogamous, only the dominant couple reproduces, once a year, and reproduction of mature subordinates is suppressed through harassment and stress.

In our study site, dominance is acquired for a duration ranging from one to 14 years. Loss of dominance is generally caused by the death of the individual during hibernation or following takeover of the territory by an individual of the same sex. During takeover, the former dominant is usually killed or, in rare cases, can reach dominance in a nearby territory (only 2 cases observed in 23 years of study). Takeover by a new dominant is followed by the infanticide of the former dominant’s litter and all same-sex subordinates are typically evicted from the territory. Marmots enter hibernation around mid-October and emerge above ground around mid-April. They soon mate and after 30 days, dominant females give birth in the main burrows. Altricial pups of ±20g are born naked, eyes closed, and digits only partially developed. The dominant females lactate for 40 days and once weaned, litters of 1 to 7 pups emerge from the natal burrow from mid-June to mid-July.

### 2.2 Study site

The study site is located in La Grande Sassìere Nature Reserve (French Alps, 45^◦^29’N, 6^◦^59’E) at an elevation of 2 400m. This is a typical alpine grassland just above the treeline presenting a high diversity of monocotyledon and dicotyledon species. The study site is a 2km long by 0.8km wide area of an alpine valley crossed by a central and permanently flowing river. Marmot territories occupy all aspects of the valley (North and South facing slopes) and population density averages 2.1 individuals.km^−2^. Chamois (*Rupicapra rupicapra*) and alpine ibex (*Capra ibex*) are the only large herbivores roaming in La Grande Sassìere Nature Reserve. Although the alpine marmot is a game species as per French law (Law of June 26th 1987), no marmots are hunted in the study site because the area is part of an officially protected area managed by the Vanoise National Park authorities.

### 2.3 Data collection

Each year since 1990 from mid-June to mid-July, between one and five observers searched territories for marmot pups using binoculars (10 × 42) and spotting scopes from 07:30am to 08:30pm. Spotting locations were in the proximity of studied territories (approx. 50m). Only territories where we expected pups were sampled. Pups were expected on a territory when the dominant female was captured pregnant or lactating and when no takeover was observed. Pups can easily be identified by their relative size, body shape and behaviour. Pups of the same litter emerge simultaneously from the same burrow. During the first 2 to 3 days, their movements are limited to this burrow but they rapidly traverse most of their territory. We assigned the same date of emergence to the whole litter. Each monitored marmot family (*k*) has, hence, one emergence date per year (*t*) noted *d_k,t_*.

On average, we monitored *N*^-^ = 23.0 families each year although this number varied between *N_t_* = 3 in 1990 and *N_t_* = 35 in 2023 depending on group dynamics and sampling effort. The latter was typically lower during the early years of the study. We qualified emergence dates according to the accuracy of the observation as some pups may have gone undetected for several days for some families because of vegetation (*e.g.* presence of fireweed *Chamaenerion angustifolium* on the burrow), difficult terrain or the split of a family leading to the emergence of unexpected litters. Most emergence dates were accurate to the day but others could be estimated with low accuracy or be completely unknown when we discovered fully active pups. Here we only considered emergence dates known with a one-day accuracy, which explains why we have less data per year than the number of monitored families. Emergence dates were unknown for only 10 families.years since 1990.

### 2.4 Statistical analyses

We coded emergence dates *d_k,t_* ∈ {1 … 54}, starting from June 1 as day 1. We analyzed emergence dates *d_k,t_* with quantile regressions (Koenker & Bassett Jr 1978). A quantile regression estimates the conditional median of the response variable in relation to predictors (see Chamailĺe-Jammes et al. (2007) for an application in ecology). We opted for this particular regression method because it is less susceptible to extreme values than the standard linear regression working on conditional means, and the method does not assume any particular probability distribution *a priori*. Another advantage of using quantile regression is the possibility to investigate changes in median date of emergence, but also changes in any quantile (*τ*) such as quartiles and deciles. If, for instance, only late emergence were to occur less and less in time, this would be detected in the last decile regression but not in the first quartile or median. In other words, quantile regression is a useful tool to explore changes in distribution of the variable of interest.

Because climate change may have stronger effects on individuals emerging early or late compared to the average, in addition to the median (*τ* = 0.50), we also explored temporal trends in the first (*τ* = 0.25) and fourth quartile (*τ* = 0.75), and in the first (*τ* = 0.10) and tenth decile (*τ* = 0.90) of the distribution of emergence dates by entering time in years (noted *T*) as an explanatory variable. The baseline model was of the following form:

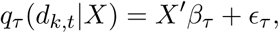

whereby *X* is the matrix of predictors (intercept and time in years *T*), *β_τ_*is the regression coefficient for quantile *τ*, and *ɛ_τ_* is the residual error variance term, which is also a function of *τ*. Since we set five different *τ* values (*τ* ∈ {0.10, 0.25, 0.50, 0.75, 0.90}), we correspondingly estimated five regression coefficients (respectively *β*_0.10_, *β*_0.25_, *β*_0.50_, *β*_0.75_ and *β*_0.90_) as well as five residual variances (respectively *ɛ*_0.10_, *ɛ*_0.25_, *ɛ*_0.50_, *ɛ*_0.75_ and *ɛ*_0.90_).

We fitted quantile regression models using the quantreg package (version 5.98; Koenker (2024)) for the R statistical software (version 4.4.1; R Core Team (2024)). We concluded that a statistically significant temporal trend occurred in emergence dates for the different quantile levels when the associated confidence limits of *β_τ_* excluded 0. To the best of our knowledge, quantile regressions do not offer standard estimation of effect sizes such as the proportion of explained variance (see Nakagawa & Cuthill 2007). We therefore estimated a pseudo-*R*^2^ by computing the correlation between observed quantile of emergence date each year and the corresponding predicted values from the quantile regression.

Note that for the sake of comparison we also ran the standard least-square regression to test for a change in mean date of pup emergence as a function of time in year. The fitted model was:

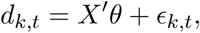

Alike quantile regression models, *X* is the matrix of predictors (intercept and time in years *T*), *θ* is the vector of regression coefficients, and *ɛ_k,t_* is a normally distributed error term.

Following Rutberg (1987) we quantified emergence synchrony from the time span, in days, covering 80% of the emergences (hence removing the earliest and latest 10% of observations). Note that if quantile regressions of the 10th and 90th percentile converge, it would mean an increase of emergence synchrony.

## 3 Results

We recorded a total of 468 emergence events of alpine marmot pups from 54 different families monitored between 1990 and 2023 at La Grande Sassìere (mean of 14.1 per year, lower than total number of monitored families because of failure to reproduce). The distribution of emergence dates (Fig. 1) was unimodal with a mean of *d*^-^ = 29.4 (between 29 and 30 June), a standard deviation of *σ_d_*= 5.7 days, and a median of *d*^∽^ = 29 (29 June). Those distribution moments suggested a slight positive skew toward late emergence dates (Pearson’s moment coefficient of skewness: *γ*_1_ = 0.31). Accordingly, the observed distribution of emergence dates was not Gaussian (Shapiro-Wilk’s normality test: *W* = 0.98, *p <* 0.001). The number of days between the 10^th^ and 90^th^ percentiles, an often used proxy of emergence synchrony (see e.g. Rutberg (1987)), was 14.0 days.

**Figure 1:**
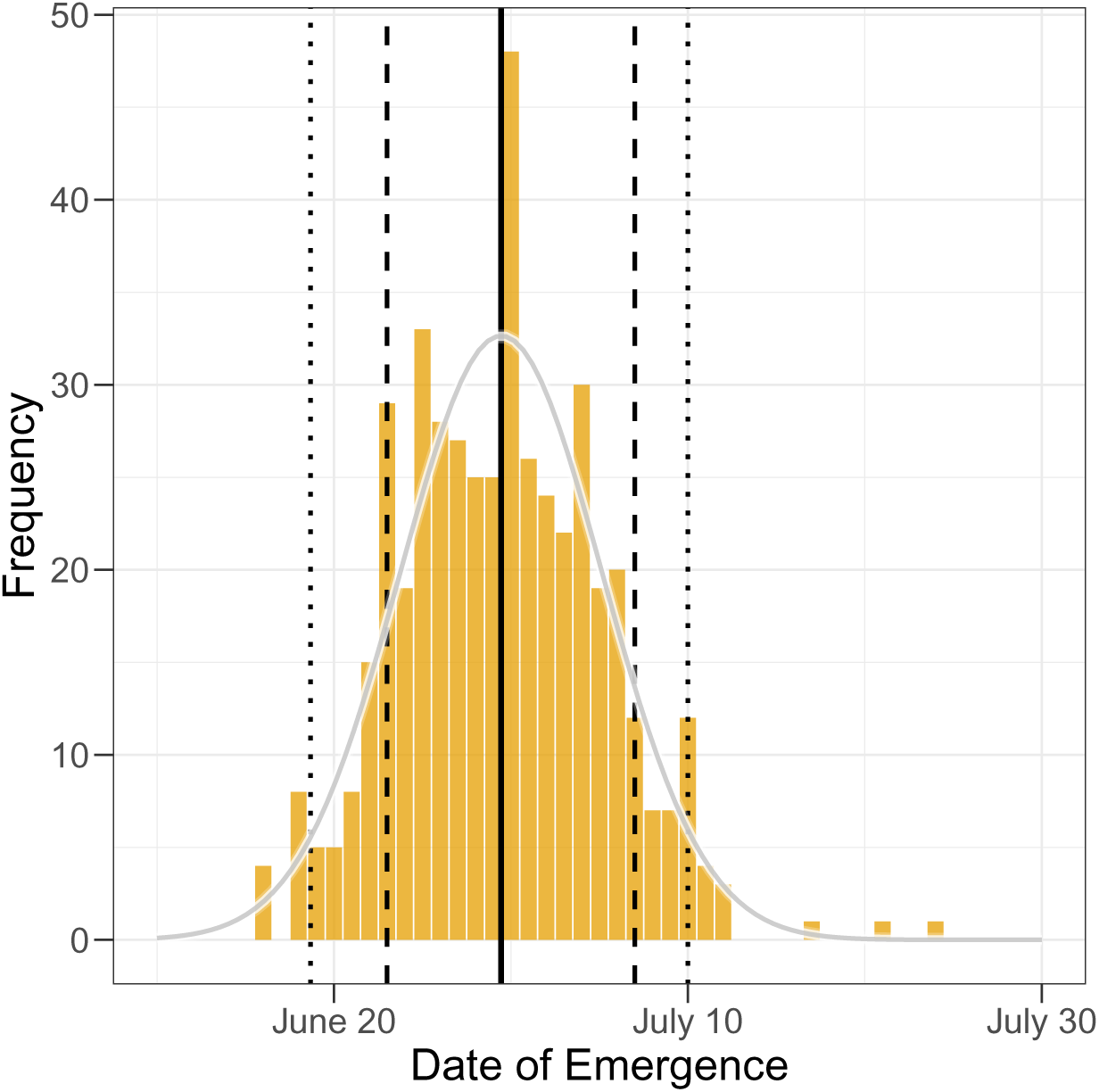
Distribution of emergence dates of marmot pups (*Marmota marmota*) observed from 1990 to 2023 at La Grande Sassìere, French Alps. Vertical lines represent distribution quantiles (0.025, 0.10, 0.50, 0.9, 0.975) and the grey continuous curve the expected values from a normal distribution taking the observed mean and variance as parameters.

The prediction of earlier dates of emergence of alpine marmot pups over time was fully supported at La Grande Sassìere (Fig. 2). All coefficients estimated by the quantile regressions were negative and statistically different from 0 (Fig. 3). The predicted advance in date of emergence was 4.7 days over 33 years for both median (*τ* = 0.50) and mean (Fig. 2) dates. The estimated coefficients varied markedly with *τ* (Fig. 3), particularly so for the first and tenth deciles (*τ* = 0.10 and 0.90), providing empirical evidence for stronger changes in time for late than early emergence events. In our study site, the 10% of the earliest emergence occurred 2.1 days earlier in 2023 than in 1990. In contrast, the 10% of latest emergence shifted by 8.2 days instead.

**Figure 2:**
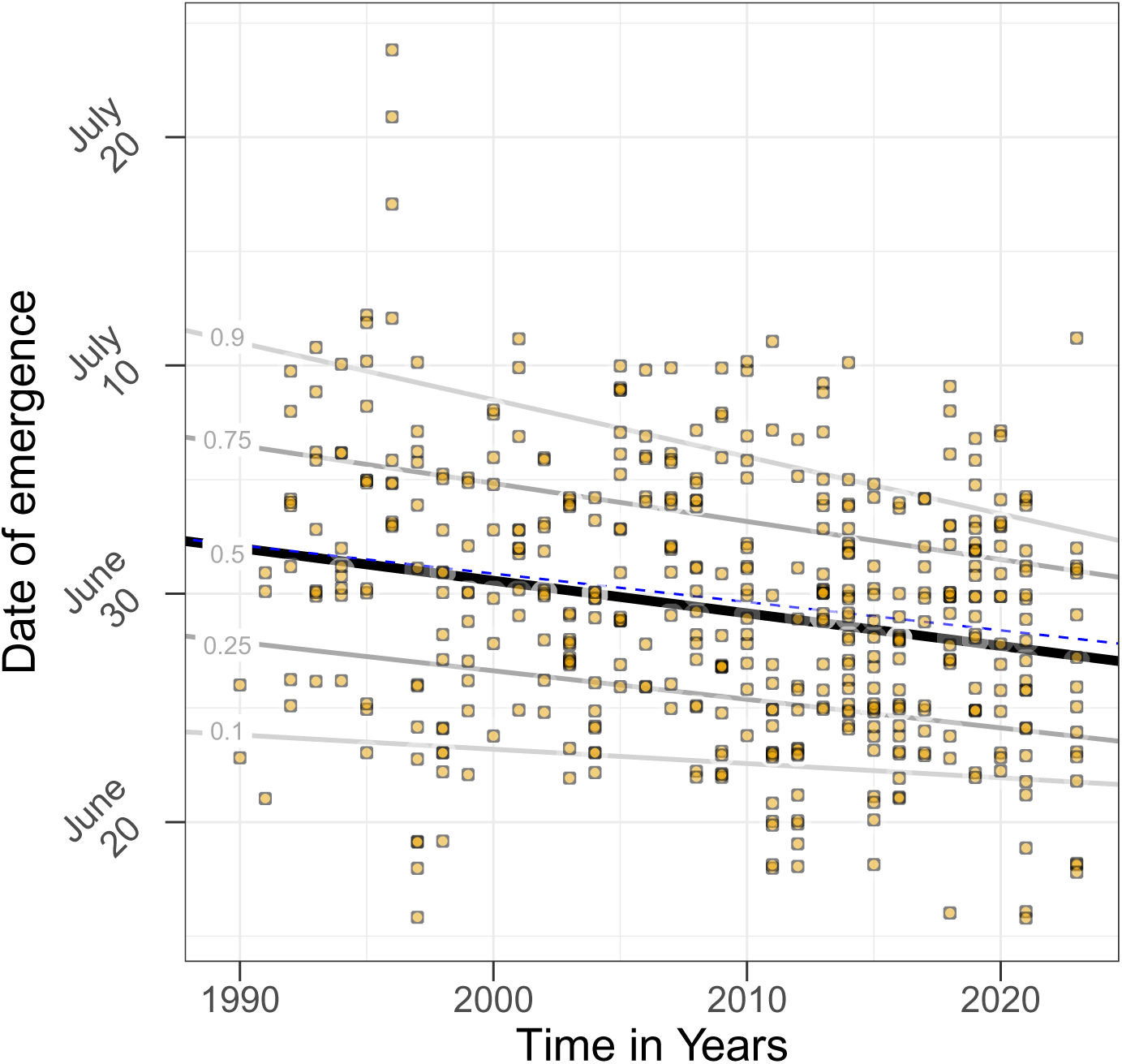
Annual variation in the emergence date of alpine marmot pups (*Marmota marmota*) observed from 1990 to 2023 at La Grande Sassìere, French Alps. We monitored a mean of 23 marmot families each year (yielding *N* = 468 emergences of pups). Overlaid are the predictions from quantile regressions whereby the black line is the predicted temporal trend of the median date of emergence, dark grey lines are the same for first and third quartiles, and light grey lines are the same for the first and last deciles (quantiles are given by the grey numbers on regression lines). Note that year 2022 is missing because of the COVID-19 field team contamination that prevented us from accessing the field site early in the season.

**Figure 3:**
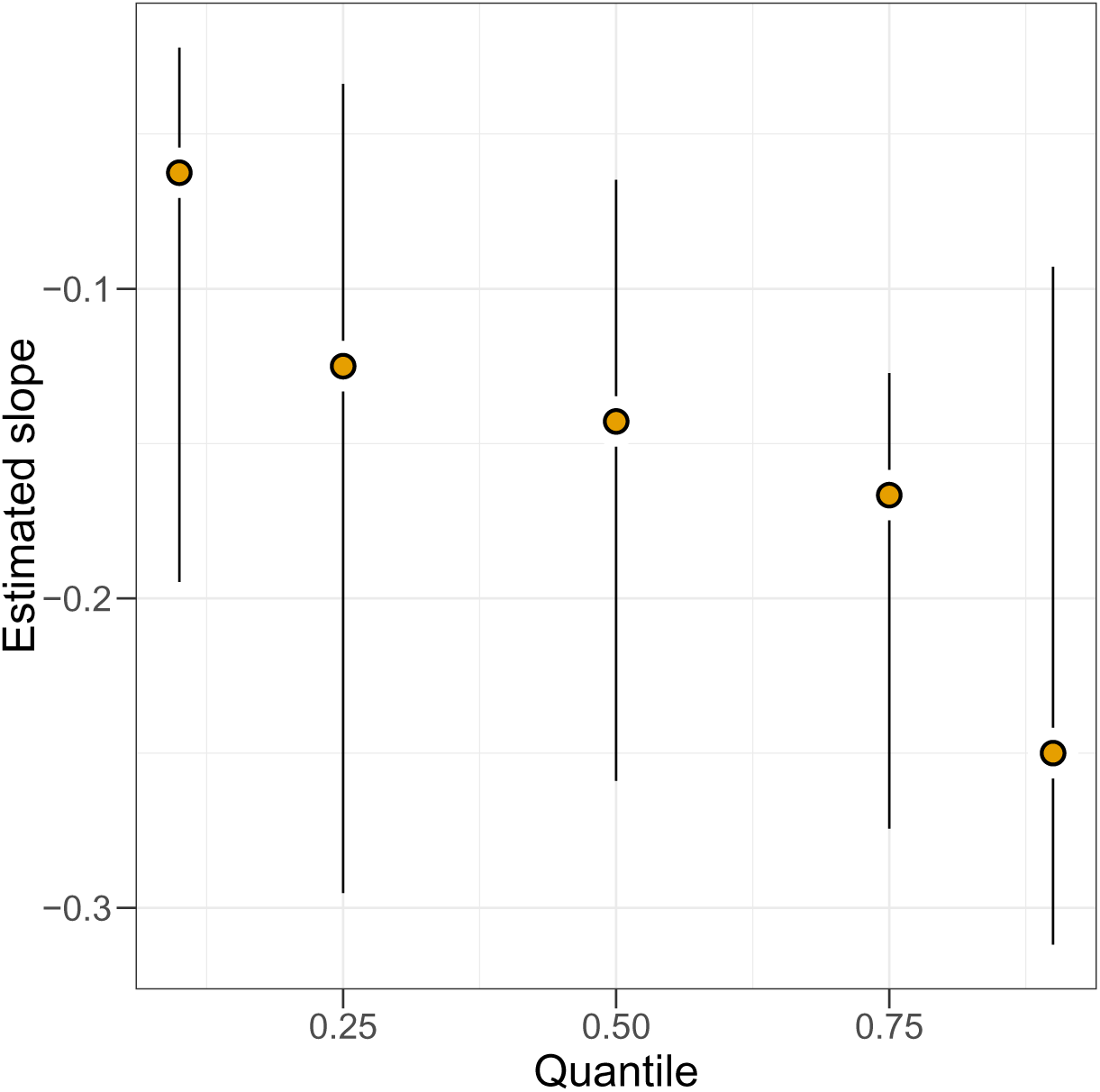
Estimated coefficients for the temporal trend estimated with a quantile regression (quantile: 0.10, 0.25, 0.50, 0.75, 0.90) on date of emergence of alpine marmot pups (*Marmota marmota*) monitored from 1990 to 2023 at La Grande Sassìere, French Alps. We monitored a mean of 23 marmot families each year for a total of *N* = 468 emergences of pups. The observed shift in phenology increases in magnitude from the earliest to the latest emergence within a reproductive season.

Although statistically significant, the proportion of variance explained by the time variable (*T*) on the median date of emergence was only 3%. Congruent with the quantile regression, a standard linear regression suggested a statistically significant decreasing time trend in mean date of pup emergence at La Grande Sassìere (Fig. 2). The estimated parameters were *θ*_1_ = _243.42_ 358.47 _473.51_ and *θ*_2_ = _−0.221_ −0.164 _−0.106_ for the model intercept and slope, respectively; the error term read *ɛ* = 5.540. This linear model accounted for *R*^2^ = 0.06 of the observed variance in emergence dates.

## 4 Discussion

The temporal variation of emergence dates of alpine marmot pups at La Grande Sassìere is no exception to the general pattern of advancing phenology (Parmesan 2006, Bronson 2009, Inouye 2022). In line with our previous observations (Tafani et al. 2013), individuals of this population track the advancing phenology of plants by showing a progressive shift of median annual pup emergence dates throughout the study period encompassing 34 years from 1990 to 2023. If the tracking of plant phenology for reproduction is supported for the alpine marmot, the prediction of a marked response of slow and hibernating species is clearly not, and more consistent with observations on the closely related yellow-bellied marmot (*Marmota flaviventris*, see Ozgul et al. 2010). The observed shift in the pup emergence phenology is, moreover, not homogeneous in the population. Litters emerging the latest are advancing about twice as fast as the earliest or average litters, leading to an increasing synchrony of pup emergence dates in time for the alpine marmot. Such a phenomenon likely emerges because of a strong time constraint for marmots between emergence from hibernation, fat accumulation, and the next hibernating event the following autumn.

With a generation time close to 7 years (ézouki 2018), the alpine marmot is a long-lived mammal relative to its specific body size, likely because of its lifestyle in terms of hibernation and sociality (Turbill et al. 2011). Recently, it has been proposed that a slow pace-of-life leads to more acute effects of climate change (Orgeret et al. 2022). This prediction was clearly not supported in the alpine marmot. The estimated rate of 0.14 days.year^−1^ we found is less than average values reported in literature (median value of 0.54 days.year^−1^ across mammalian populations, Table 1). The estimated change is 2.8 times smaller to that observed in the alpine chamois living close to our study area (Kourkgy et al. 2016). More similar are pups of the closely related yellow-bellied marmot that emerge increasingly earlier at a rate of −0.17 days.year^−1^ (Ozgul et al. 2010). Overall, the magnitude in the shift we report in pup emergence phenology is of moderate magnitude given the life history and pace-of-life of the alpine marmot. The only exception is the roe deer (*Capreolus capreolus*) whose mean birth dates show little to no advance with climate change (rate of −0.08 days.year^−1^; Table 1). That roe deer reproductive cycle depends mostly on day length variation rather than temperature (Plard et al. 2014) makes this species unusual among mammals.

Our result on alpine marmot rather suggest a lesser sensibility of hibernating species phenology to climate changes. Accordingly, a comparative study stresses that hibernating species are less likely to be endangered because of their relatively lower exposure to environmental stress such as climate changes compared to non-hibernators (Liow et al. 2009). Another mechanism is that hibernation could constrain the ability of alpine marmot to track plant phenology and the onset of spring. The approximate time-lag between mating and pup emergence, the variable we recorded and analyzed, is two and a half months. Mating usually takes place right after the end of hibernation (Müller-Using 1957) followed by a one-month gestation (between 33 and 35 days, Psenner 1957). Newborns then grow underground for 40 days after which they will emerge from the burrow for the first time (Wilson et al. 2016). We do not know whether earlier emergence dates of alpine marmot pups over time are due to earlier parturition date, shorter lactation period, or both. Average spring emergence date of adult yellow-bellied marmots also is advancing (Inouye et al. 2000). Consequently, pup emergence date could indirectly correlate with the timing of hibernation of mothers, and particularly the end of it.

Since fossorial sciurids hibernate in the dark, environmental cues outside of the burrow (photoperiod, temperature, food availability) have little influence making the role of the circannual clock predominant. In addition, little is known yet on the proximate causes determining the end of torpor bouts (heterothermic phase Williams et al. 2012). The timing of hibernation seems primarily governed by an endogeneous circannual clock that needs external cues to maintain synchrony with seasons (Heller & Poulson 1970). Photoperiod is likely involved in the calibration of circannual rythms (Ikegami & Yoshimura 2012), which will not change with climate change (see the case of roe deer, Plard et al. 2014, for similar argument). If very little is known on the physiology of the circannual clock, a few studies nevertheless have shown an association between emergence date and mean temperature (Wells et al. 2022). Although the physiological pathway linking air temperature and emergence from hibernation is not well understood yet, it could indirectly account for the temporal trends of pup emergence in the alpine marmot.

An alternative could be a matter of energy consumption while hibernating, and reduced body fat accumulation of females during spring and summer. At La Grande Sassìere, females are now in poorer body condition than in the 1990s when the monitoring started (Tafani et al. 2013, Canale et al. 2016). In species with short-term torpor, individuals may not enter in torpor if their energy reserve is too limited to ensure the costly full rewarming of their body (Galster & Morrison 1975). The same could happen in alpine marmots, whereby hibernation duration could be shortened by one or two bouts of low temperature because they started hibernating in poorer condition in the preceding autumn. The energy shortage hypothesis would predict an association between autumn body condition of individuals and date of emergence in spring. For instance, a correlation between autumn snow cover and date of emergence from hibernation was reported for Arctic ground squirrels (*Urocitellus parryii,* Sheriff et al. 2015).

Besides the general shift in average timing of pup emergence we report for alpine marmots over the last 30 years, climate change seemingly affects the length of the birth season too, as previously documented on reindeer *Rangifer tarandus* in Scandinavia (Paoli et al. 2018). While the predicted span of emergence dates was 17 days in the 90ies, 80% of pup emergences occurred within 13 days 30 years later at La Grande Sassìere (−26%). The range of birth dates, also referred to *birth synchrony* (see Thel et al. 2022), is an important characteristic of birth phenology which, like the timing, should be under selective forces (Sinclair et al. 2000). Two non-mutually exclusive hypotheses may explain this marked increase in birth synchrony in the alpine marmots.

First, the predator swamping hypothesis posits that high synchrony of births of prey species would limit predation by diluting risks among conspecifics (Rutberg 1987, Sinclair et al. 2000). Two lines of evidence give little support to this mechanism in shaping the higher birth synchrony we report for alpine marmots. The main predador of alpine marmots, the golden eagle (*Aquila chrysaetos*), has always been present in the Alps as has been the red fox (*Vulpes vulpes*) been too. Wolf (*Canis lupus*) that recently returned in our study site (with first observations in 2018-2019) seldom prey upon marmots, and cannot account for the progressive temporal trend we observed in emergence phenology of pups. Second, if at work, this hypothesis cannot explain the stronger temporal change of late compared to early pup emergences, because individuals born at both ends of the distribution would face similar high predation risks. A normalizing selection would be rather expected under this hypothesis.

An alternative explanation accounting for the higher synchrony of pup emergences would be a directional selection against late litters. In seasonal environment, individuals born late rapidly face unfavourable early-life conditions, and usually suffer from higher juvenile mortality than those born earlier (Plard et al. 2014). For the alpine marmot, pup emerging relatively late in a reproductive season will have limited body growth before food resource quantity and quality start to decrease in summer (Paterson et al. 2022). Like most mammals, light individuals are more at risk to die from predation or starvation and could be counter-selected on the long run. In the context of climate change, the life history consequences of a given time-lag between emergence date and spring likely differs between the earliest and latest pups (Miller-Rushing et al. 2010). During a reproductive season, late-born individuals lag behind plant phenology to a larger extent than early-born individuals because plant phenology advance faster than their consumers (Iler et al. 2021). A spring occurring 1-day earlier will lead to a disproportionate relative lag for early compared to late emergences, with similar consequence for body growth. This phenomenon is known as the multiplier effect and arises because body growth is a non-linear process (White 1983). Such an asymmetry of selective forces could also lead to directional selection and, hence, to higher synchrony of births and emergences at the population level. The fraction of the population born early may in turn show relatively weaker changes in phenology compared to the late-born individuals.

Our results are broadly consistent with the expectation of a shift toward earlier phenology of reproduction and emergence of pups in the alpine marmot. Even if the data analyses based on quantile regressions are robust to extreme values (*i.e.* very early or very late emergence), data collection methodology may have affected the temporal pattern of emergence synchrony we report. By design, we could not detect and record very late litter emergence after the field season ended in late July. Missed litters are, however, easily detectable at the next field season by observations of yearlings in a territory with no record of pups the year before – assuming no complete loss of the litter the first winter. Missed litters occurred in *<* 1% of all records (4/491), with no evidence of a time trend in its occurrence (1995, 1996, 2005 and 2014), hence suggesting very limited bias in our data.

## Acknowledgments

We thank Sylvia Pardonnet, Jeanne Duhayer and all volunteering fieldworkers for their invaluable contribution to data collection. We also extend our acknowledgments to the municipality of Tignes for their long-term logistical support to the marmot project. This work has been financially supported by the Agence National de la Recherche (ANR SOCIALIPOP) and the SEE-Life initiative from the INEÉE-CNRS. Our thanks are extended to two anonymous reviewers who helped at improving the clarity and writing of our manuscript.

## Notes

### Competing Interest Statement

The authors have declared no competing interest.

### Summary of Updates

Data, statistics and clarification of text

## References

Abrahms, B., Rafiq, K., Jordan, N. R. & McNutt, J. (2022), ‘Long-term, climate-driven phenological shift in a tropical large carnivore’, Proceedings of the National Academy of Sciences 119(27), e2121667119.

Adamík, P. & Kŕal, M. (2008), ‘Climate-and resource-driven long-term changes in dormice populations negatively affect hole-nesting songbirds’, Journal of Zoology 275(3), 209–215.

Armitage, K. B. (2014), Marmot biology: sociality, individual fitness, and population dynamics, Cambridge University Press.

Armitage, K. B. (2017), ‘Hibernation as a major determinant of life-history traits in marmots’, Journal of Mammalogy 98(2), 321–331.

Badeck, F.-W., Bondeau, A., Böttcher, K., Doktor, D., Lucht, W., Schaber, J. & Sitch, S. (2004), ‘Responses of spring phenology to climate change’, New phytologist 162(2), 295–309.

Beniston, M. (2006), ‘Mountain weather and climate: a general overview and a focus on climatic change in the Alps’, Hydrobiologia 562, 3–16.

Bonnet, T., Morrissey, M. B., Morris, A., Morris, S., Clutton-Brock, T. H., Pemberton, J. M. & Kruuk, L. E. (2019), ‘The role of selection and evolution in changing parturition date in a red deer population’, PLoS Biology 17(11), e3000493.

Boutin, S. & Lane, J. E. (2014), ‘Climate change and mammals: evolutionary versus plastic responses’, Evolutionary applications 7(1), 29–41.

Bronson, F. (2009), ‘Climate change and seasonal reproduction in mammals’, Philosophical Transactions of the Royal Society B: Biological Sciences 364(1534), 3331– 3340.

Bronson, F. H. (1989), Mammalian reproductive biology, University of Chicago Press.

Bunnell, F. L. (1982), ‘The lambing period of mountain sheep: synthesis, hypotheses, and tests’, Canadian Journal of Zoology 60(1), 1–14.

Burthe, S., Butler, A., Searle, K. R., Hall, S. J., Thackeray, S. J. & Wanless, S. (2011), ‘Demographic consequences of increased winter births in a large aseasonally breeding mammal (*Bos taurus*) in response to climate change’, Journal of Animal Ecology 80(6), 1134–1144.

Canale, C. I., Ozgul, A., Allaińe, D. & Cohas, A. (2016), ‘Differential plasticity of size and mass to environmental change in a hibernating mammal’, Global Change Biology 22(10), 3286–3303.

Chamailĺe-Jammes, S., Fritz, H. & Murindagomo, F. (2007), ‘Detecting climate changes of concern in highly variable environments: Quantile regressions reveal that droughts worsen in Hwange National Park, Zimbabwe’, Journal of Arid Environments 71(3), 321–326.

Cohen, J. M., Lajeunesse, M. J. & Rohr, J. R. (2018), ‘A global synthesis of animal phenological responses to climate change’, Nature Climate Change 8(3), 224–228.

Cushing, D. (1973), ‘The natural regulation of fish populations’, Sea fisheries research pp. 399–411.

Davis, D. E. (1976), ‘Hibernation and circannual rhythms of food consumption in marmots and ground squirrels’, The Quarterly Review of Biology 51(4), 477–514.

Dobson, F. S. (2007), ‘A lifestyle view of life-history evolution’, Proceedings of the National Academy of Sciences 104(45), 17565–17566.

Galster, W. & Morrison, P. (1975), ‘Gluconeogenesis in arctic ground squirrels between periods of hibernation’, American Journal of Physiology-Legacy Content 228(1), 325–330.

Gamelon, M., Besnard, A., Gaillard, J.-M., Servanty, S., Baubet, E., Brandt, S. & Gimenez, O. (2011), ‘High hunting pressure selects for earlier birth date: wild boar as a case study’, Evolution 65(11), 3100–3112.

Geiser, F. (2013), ‘Hibernation’, Current Biology 23(5), R188–R193.

Gienapp, P., Teplitsky, C., Alho, J., Mills, J. & Merilä, J. (2008), ‘Climate change and evolution: disentangling environmental and genetic responses’, Molecular ecology 17(1), 167–178.

Hagen, R., Ortmann, S., Elliger, A. & Arnold, J. (2021), ‘Advanced roe deer (capreolus capreolus) parturition date in response to climate change’, Ecosphere 12(11), e03819.

Heller, H. C. & Poulson, T. L. (1970), ‘Circanian rhythms—II. endogenous and exogenous factors controlling reproduction and hibernation in chipmunks (Eutamias) and ground squirrels (Spermophilus)’, Comparative Biochemistry and Physiology 33(2), 357–383.

Hock, R., Rasul, G., Adler, C., Ćaceres, B., Gruber, S., Hirabayashi, Y., Jackson, M., Kääb, A., Kang, S., Kutuzov, S., Milner, A., Molau, U., Morin, S., Orlove, B. & Steltzer, H. (2019), High mountain areas, in H.-O. Pörtner, D. Roberts, V. Masson-Delmotte, P. Zhai, M. Tignor, E. Poloczanska, K. Minten-beck, A. Alegŕıa, M. Nicolai, A. Okem, J. Petzold, B. Rama & W. N.M., eds, ‘IPCC special report on the ocean and cryosphere in a changing climate’, Cambridge University Press, Cambridge, UK and New York, NY, USA, pp. 131–202.

Ikegami, K. & Yoshimura, T. (2012), ‘Circadian clocks and the measurement of daylength in seasonal reproduction’, Molecular and cellular endocrinology 349(1), 76–81.

Iler, A. M., CaraDonna, P. J., Forrest, J. R. & Post, E. (2021), ‘Demographic consequences of phenological shifts in response to climate change’, Annual Review of Ecology, Evolution, and Systematics 52(1), 221–245.

Inouye, D. W. (2022), ‘Climate change and phenology’, Wiley Interdisciplinary Reviews: Climate Change 13(3), e764.

Inouye, D. W., Barr, B., Armitage, K. B. & Inouye, B. D. (2000), ‘Climate change is affecting altitudinal migrants and hibernating species’, Proceedings of the National Academy of Sciences 97(4), 1630–1633.

Inouye, D. W. & Wielgolaski, F. E. (2025), Phenology at high altitudes, in ‘Phenology: An integrative environmental science’, Springer, pp. 281–311.

Koenker, R. (2024), quantreg: Quantile Regression. R package version 5.98. **URL:** https://CRAN.R-project.org/package=quantreg

Koenker, R. & Bassett Jr, G. (1978), ‘Regression quantiles’, Econometrica: journal of the Econometric Society pp. 33–50.

Kourkgy, C., Garel, M., Appolinaire, J., Loison, A. & Töıgo, C. (2016), ‘Onset of autumn shapes the timing of birth in Pyrenean chamois more than onset of spring’, Journal of Animal Ecology 85(2), 581–590.

Lane, J. E., Kruuk, L. E., Charmantier, A., Murie, J. O. & Dobson, F. S. (2012), ‘Delayed phenology and reduced fitness associated with climate change in a wild hibernator’, Nature 489(7417), 554–557.

Lenoir, J., Gégout, J.-C., Marquet, P. A., de Ruffray, P. & Brisse, H. (2008), ‘A significant upward shift in plant species optimum elevation during the 20th century’, science 320(5884), 1768–1771.

Liow, L. H., Fortelius, M., Lintulaakso, K., Mannila, H. & Stenseth, N. C. (2009), ‘Lower extinction risk in sleep-or-hide mammals’, The American Naturalist 173(2), 264–272.

Loewen, C. J., Jackson, D. A. & Gilbert, B. (2023), ‘Biodiversity patterns diverge along geographic temperature gradients’, Global Change Biology 29(3), 603–617.

Mahoney, P. J., Joly, K., Borg, B. L., Sorum, M. S., Rinaldi, T. A., Saalfeld, D., Golden, H., Latham, A. D. M., Kelly, A. P., Mangipane, B. et al. (2020), ‘Denning phenology and reproductive success of wolves in response to climate signals’, Environmental Research Letters 15(12), 125001.

Menzel, A., Sparks, T. H., Estrella, N., Koch, E., Aasa, A., Ahas, R., Alm-Kübler, K., Bissolli, P., Braslavská, O., Briede, A., et al. (2006), ‘European phenological response to climate change matches the warming pattern’, Global change biology 12(10), 1969–1976.

Michener, G. (1984), Age, sex, and species differences in the annual cycles of ground-dwelling sciurids: implications for sociality, in ‘The Biology of Ground Dwelling Squirrels: Annual Cycles, Behavioral Ecology and Sociality’, University of Nebraska Press, pp. 81–107.

Miller-Rushing, A. J., Høye, T. T., Inouye, D. W. & Post, E. (2010), ‘The effects of phenological mismatches on demography’, Philosophical Transactions of the Royal Society B: Biological Sciences 365(1555), 3177–3186.

Monahan, W. B., Rosemartin, A., Gerst, K. L., Fisichelli, N. A., Ault, T., Schwartz, M. D., Gross, J. E. & Weltzin, J. F. (2016), ‘Climate change is advancing spring onset across the US national park system’, Ecosphere 7(10), e01465.

Morren, C. (1849), ‘Le globe, le temps et la vie’, Bulletins de l’Académie royale des Sciences, des Lettres et des Beaux-Arts de Belgique 16(2), 660–684.

Moyes, K., Nussey, D. H., Clements, M. N., Guinness, F. E., Morris, A., Morris, S., Pemberton, J. M., Kruuk, L. E. & Clutton-Brock, T. H. (2011), ‘Advancing breeding phenology in response to environmental change in a wild red deer population’, Global change biology 17(7), 2455–2469.

Müller-Using, D. (1957), ‘Die Paarungsbiologie des Murmeltieres’, Zeitschrift fur Jagdwissenschaft 3(1), 24–28.

Nakagawa, S. & Cuthill, I. C. (2007), ‘Effect size, confidence interval and statistical significance: a practical guide for biologists’, Biological reviews 82(4), 591–605.

Orgeret, F., Thiebault, A., Kovacs, K. M., Lydersen, C., Hindell, M. A., Thompson, S. A., Sydeman, W. J. & Pistorius, P. A. (2022), ‘Climate change impacts on seabirds and marine mammals: The importance of study duration, thermal tolerance and generation time’, Ecology Letters 25(1), 218–239.

Osinga, N., Pen, I., De Haes, H. U. & Brakefield, P. M. (2012), ‘Evidence for a progressively earlier pupping season of the common seal (*Phoca vitulina*) in the Wadden Sea’, Journal of the Marine Biological Association of the United Kingdom 92(8), 1663–1668.

Ozgul, A., Childs, D. Z., Oli, M. K., Armitage, K. B., Blumstein, D. T., Olson, L. E., Tuljapurkar, S. & Coulson, T. (2010), ‘Coupled dynamics of body mass and population growth in response to environmental change’, Nature 466(7305), 482– 485.

Paoli, A., Weladji, R. B., Holand, Ø. & Kumpula, J. (2018), ‘Winter and spring climatic conditions influence timing and synchrony of calving in reindeer’, PloS one 13(4), e0195603.

Parmesan, C. (2006), ‘Ecological and evolutionary responses to recent climate change’, *Annual Reviews of Ecology*, Evolution and Systematics 37(1), 637–669.

Parmesan, C. & Yohe, G. (2003), ‘A globally coherent fingerprint of climate change impacts across natural systems’, nature 421(6918), 37–42.

Paterson, J. T., Proffitt, K. M., DeCesare, N. J., Gude, J. A. & Hebblewhite, M. (2022), ‘Evaluating the summer landscapes of predation risk and forage quality for elk (Cervus canadensis)’, Ecology and evolution 12(8), e9201.

Plard, F., Gaillard, J.-M., Coulson, T., Hewison, A. M., Delorme, D., Warnant, C. & Bonenfant, C. (2014), ‘Mismatch between birth date and vegetation phenology slows the demography of roe deer’, PLoS biology 12(4), e1001828.

Post, E. & Forchhammer, M. C. (2008), ‘Climate change reduces reproductive success of an arctic herbivore through trophic mismatch’, Philosophical transactions of the Royal Society B: Biological sciences 363(1501), 2367–2373.

Psenner, H. (1957), ‘Neues vom Murmeltier, Marmota m. marmota (Linne, 1758)’, Säugetierkundliche Mitteilungen 5, 4–10.

R Core Team (2024), R: A Language and Environment for Statistical Computing, R Foundation for Statistical Computing, Vienna, Austria. URL: https://www.R-project.org/

Radchuk, V., Reed, T., Teplitsky, C., van de Pol, M., Charmantier, A., Hassall, C., Adamík, P., Adriaensen, F., Ahola, M. P., Arcese, P., et al. (2019), ‘Adaptive responses of animals to climate change are most likely insufficient’, Nature communications 10(1), 3109.

éale, D., McAdam, A. G., Boutin, S. & Berteaux, D. (2003), ‘Genetic and plastic responses of a northern mammal to climate change’, Proceedings of the Royal Society of London. Series B: Biological Sciences 270(1515), 591–596.

Rehnus, M., Peĺaez, M. & Bollmann, K. (2020), ‘Advancing plant phenology causes an increasing trophic mismatch in an income breeder across a wide elevational range’, Ecosphere 11(6), e03144.

ézouki, C. (2018), The influence of lifestyle on demographic responses to climate change: the Alpine marmot as a case study, PhD thesis, Université de Lyon.

ézouki, C., Tafani, M., Cohas, A., Loison, A., Gaillard, J.-M., Allaińe, D. & Bonenfant, C. (2016), ‘Socially mediated effects of climate change decrease survival of hibernating alpine marmots’, Journal of Animal Ecology 85(3), 761–773.

Rutberg, A. T. (1987), ‘Adaptive hypotheses of birth synchrony in ruminants: an interspecific test’, The American Naturalist 130(5), 692–710.

Sheriff, M. J., Buck, C. L. & Barnes, B. M. (2015), ‘Autumn conditions as a driver of spring phenology in a free-living arctic mammal’, Climate Change Responses 2, 1–8.

Sinclair, A., Mduma, S. A. & Arcese, P. (2000), ‘What determines phenology and synchrony of ungulate breeding in Serengeti?’, Ecology 81(8), 2100–2111.

Stenseth, N. C. & Mysterud, A. (2002), ‘Climate, changing phenology, and other life history traits: nonlinearity and match–mismatch to the environment’, Proceedings of the National Academy of Sciences 99(21), 13379–13381.

Tafani, M., Cohas, A., Bonenfant, C., Gaillard, J.-M. & Allaińe, D. (2013), ‘Decreasing litter size of marmots over time: a life history response to climate change?’, Ecology 94(3), 580–586.

Thel, L., Chamailĺe-Jammes, S. & Bonenfant, C. (2022), ‘How to describe and measure phenology? an investigation on the diversity of metrics using phenology of births in large herbivores’, Oikos 2022(4), e08917.

Turbill, C., Bieber, C. & Ruf, T. (2011), ‘Hibernation is associated with increased survival and the evolution of slow life histories among mammals’, Proceedings of the Royal Society B: Biological Sciences 278(1723), 3355–3363.

Visser, M. E. & Both, C. (2005), ‘Shifts in phenology due to global climate change: the need for a yardstick’, Proceedings of the Royal Society B: Biological Sciences 272(1581), 2561–2569.

Wells, C. P., Barbier, R., Nelson, S., Kanaziz, R. & Aubry, L. M. (2022), ‘Life history consequences of climate change in hibernating mammals: a review’, Ecography 2022(6), e06056.

White, R. G. (1983), ‘Foraging patterns and their multiplier effects on productivity of northern ungulates’, Oikos 40, 377–384.

Wielgolaski, F. E. & Inouye, D. W. (2003), ‘High latitude climates’, Phenology: An integrative environmental science pp. 175–194.

Williams, C. T., Barnes, B. M., Richter, M. & Buck, C. L. (2012), ‘Hibernation and circadian rhythms of body temperature in free-living arctic ground squirrels’, Physiological and Biochemical Zoology 85(4), 397–404.

Wilson, D. E., Lacher Jr, T. E. & Mittermeier, R. A. (2016), Handbook of the mammals of the world. Volume 6: Lagomorphs and Rodents I, Lynx Edicions.

